# Anti-snake behavior and snake discrimination in vervet monkeys

**DOI:** 10.1101/2024.07.04.602064

**Authors:** Lukas Schad, Erica van de Waal, Julia Fischer

## Abstract

Anti-predator behavior allows to investigate how animals classify potential threats in their environment and which cognitive mechanisms might be involved in risk assessment. Snakes are common predators for many primate species, yet most of our knowledge on primate anti-snake behavior stems from predator model experiments. Few studies have investigated natural predator-prey interactions. Here, we combine an observational study on anti-snake behavior in free-ranging vervet monkeys, *Chlorocebus pygerythrus,* with an experimental study of snake classification, to achieve an integrated understanding of the monkeys’ responses to and classification of snakes. Over 25 months, we gathered data from over 200 individuals in five groups and recorded natural encounters between vervet monkeys and five different species of snakes. We assessed whether the time that monkeys spent inspecting snakes from close by varied with the encountered species. We further examined whether age and sex classes differed in their propensity to inspect snakes or alarm call. Pythons were associated with higher average inspection durations and were more likely to elicit inspection responses. While adult males were less likely to approach and inspect snakes, we found no effect of age or sex on alarm calling probability. Alarm calling appeared to be restricted to individuals in close proximity to snakes, suggesting that recruitment of nearby group members is an essential function of this call type. In the experiments, we tested whether vervet monkeys discriminated snake models by length and/or diameter, but found no effect of model size on the intensity of anti-snake behavior. While the experimental data showed the same trends as data from natural encounters, responses towards model pythons were much stronger than responses towards real pythons. These results point to a potential sampling bias in predator model experiments that needs to be taken into account when assessing data from experiments alone.

## 1 | Introduction

Snakes are ancestral predators of mammals and many species that face snake predation respond with anti-predator behaviors like mobbing and alarm calling (Coss et al., 2007; Crofoot, 2012; Hernández Tienda et al., 2021; Isbell, 2006; León et al., 2023; Mielke et al., 2019; Price & Fischer, 2014; Ramakrishnan et al., 2005; Range & Fischer, 2004; Seyfarth et al., 1980b; Weiss et al., 2015; Wheeler, 2008). Anti-predator behavior has been studied in great detail as it can inform our understanding of the function of warning signals, the mechanisms guiding signal usage and comprehension, as well as the cognitive processes underlying predator recognition and classification (Caro, 2005; Fischer, 2022; Maynard Smith & Harper, 2003; Zuberbühler, 2009).

Since data on natural predator-prey interactions are difficult to obtain, predator models have become a common tool to simulate the presence of a threat experimentally (Fischer, 2022). Snake models have been used to study whether non-human primates (hereafter ‘primates’) modify call production depending on the knowledge state of their audience (Crockford et al., 2012, 2017; Girard-Buttoz et al., 2020; León et al., 2023; Mielke et al., 2019; Schel et al., 2013) and to examine the mechanisms that underpin rapid predator detection in primates (Etting et al., 2014; Isbell, 2006; Isbell & Etting, 2017; Masataka et al., 2018; Ramakrishnan et al., 2005; Van Le et al., 2014; Wheeler et al., 2011; Zeller, Ballesta, et al., 2023; Zeller, Garcia, et al., 2023).

However, there is a lack of systematic observational studies investigating prey responses to real snakes in their natural habitats, especially in the primate literature. Although such studies are essential to pin down the drivers of anti-snake behavior, non-experimental accounts of primate snake interactions are mostly limited to non-systematic case studies or they have a low sample sizes (Bartecki & Heymann, 1987; Chapman, 1986; Digweed et al., 2005; Ferrari & Beltrão-Mendes, 2011; Ferrari & Ferrari, 1990; Gursky, 2006; Perry et al., 2003; Srivastava, 1991). As a result, we still know comparatively little about the function and mechanism of primate anti-snake behavior. There is some indirect support for kin selection as a potential driver of snake alarm calling in capuchin monkeys, since immigrant males were found to exhibit lower alarm calling probabilities than philopatric females (Wheeler, 2008). In some species of ground dwelling rodents, females with dependent offspring were also reported to display higher rates of mobbing and alarm calling, indicating a role of kin selection (Phillips & Waterman, 2014; Swaisgood et al., 1999, 2003). Mobbing of snakes has also been argued to serve as a form of perception advertisement (Carlson & Griesser, 2022; Crofoot, 2012; Curio, 1978), incentivizing snakes to relocate to different hunting grounds (Clark, 2005; Gursky, 2006; Swaisgood et al., 1999).

Many primates encounter a range of different snake species, some of which actively prey on them, while others are not genuine predators but may still represent deadly hazards in case of venomous snakes. While classifying potential predators into aerial, terrestrial and snake-like threats might be innate (Fischer & Price, 2017; Seyfarth & Cheney, 1980, 1986; Wegdell et al., 2019), fine tuning of behavioral responses and discriminating between threats that belong to the same predator class, e.g. different snake species, appears to be guided by experience and potentially social learning (Coss & Biardi, 1997; León et al., 2023; Meno et al., 2013; Owings et al., 2001; Putman et al., 2015; Seyfarth & Cheney, 1986; Srivastava, 1991; Towers & Coss, 1990). A few studies have addressed the question whether primates discriminate snakes species according to the risk they pose (Hernández Tienda et al., 2021; León et al., 2023; Meno et al., 2013; Ouattara et al., 2009; Ramakrishnan et al., 2005). Potential snake features that have been proposed to affect prey responses include snake size, scale patterning, head shape and posture (Etting & Isbell, 2014; Hernández Tienda et al., 2021; Isbell & Etting, 2017; Meno et al., 2013; Ramakrishnan et al., 2005; Swaisgood et al., 1999). Whether primates discriminate among different snake species and by which mechanisms this discrimination is achieved, can thus inform our conceptualization of the cognitive processes that guide predator classification and risk assessment in primates (Fischer, 2022).

Vervet monkeys (*Chlorocebus pygerythrus*) lend themselves to investigating anti-snake behavior, as this species has been under immense predation pressure, also evident in their alarm call system (Struhsaker, 1967). In response to their main predator classes, including mammalian carnivores, raptors and snakes, vervet monkeys respond with acoustically distinct alarm calls (Seyfarth et al., 1980b, 1980a). Upon detecting a snake, vervet monkeys typically assume a bipedal stance and produce a characteristic alarm call also known as ‘chutter’ (Struhsaker, 1967). The acoustic structure of this call likely facilitates localizing the signaler (Digweed et al., 2005; Marler, 1955; Struhsaker, 1967), and other individuals tend to respond to it by assuming a bipedal posture themselves and scanning the ground (Cheney & Seyfarth, 1990; Seyfarth et al., 1980a, 1980b).

Vervet monkeys live in multi-male multi-female groups with female philopatry and are seasonal breeders exhibiting female codominance and moderate male reproductive skew (Bonnell et al., 2021; Hemelrijk et al., 2020; Minkner et al., 2018; Saccà et al., 2022; Young et al., 2017). Around the time of sexual maturity, males disperse from their natal groups and thereafter keep migrating among groups with an average residency length of about two mating seasons (Henzi & Lucas, 1980; Young et al., 2019). As the dispersing sex, adult males can be expected to have fewer kin in the group relative to females and since average male tenure does not exceed two mating seasons, most males will only have a limited number of offspring in their present group.

To characterize the function of anti-snake behavior, we examined how frequently individuals of different age and sex groups inspected snakes within close proximity and whether they produced alarms calls in snake encounters. If kin selection would be a driving force of anti-snake behavior, males might, on average, be expected to profit less from alarm calling than other age and sex groups (Wheeler, 2008). Pythons are the only confirmed snake predators of vervets, both in general and in our population, but bites from venomous snakes have been reported as a cause of mortality in other populations (Ducheminsky et al., 2014; Henzi et al., 2021). If vervets would attribute varying degrees of risk to the different snake species they encounter, this classification should affect their behavior. We therefore investigated whether the likelihood that individuals inspected snakes or alarm called in encounters varied with the encountered snake species. We further examined how long groups monitored the different snake species from close by, to determine whether species evoked different responses on a group level.

While pythons are distinct from other encountered snake species in various morphological attributes, their length and diameter tends to exceed that of other species in general. An efficient heuristic for discriminating predatory from non-predatory snakes would thus be to classify snakes by size (Ramakrishnan et al., 2005). In the second part of the study, we therefore presented python models of different length and diameter to investigate whether vervet monkeys classify snakes by size. We expected that monkeys would attribute a higher risk to larger snake models and therefore predicted that an increase in model length and diameter would be associated with higher alarm calling and inspection probability.

## 2 | Methods

### 2.1 | Study site and subjects

Data were collected from six habituated groups of free-ranging Vervet monkeys, between April 2020 and April 2022, at the INKAWU Vervet Project, Mawana Game Reserve (28°00.327S, 031°12.348E), KwaZulu-Natal, South Africa. Over the 25 month study period, group composition naturally varied due to mortality, birth, maturation, dispersal and immigration. All subjects (monkeys with age ≥ 6 months) could be identified individually via morphological features. Data could not be recorded blindly because our study required observers recognized animals in the field. From the six study groups (ESM, Table 1), one group (CR) was re-habituated in early 2021 after it had been without observers for most of 2020 due to the COVID-19 pandemic, resulting in comparatively few observations.

### 2.2 | Reptile species

The most frequently encountered snakes that elicit alarm calls from monkeys in our population are the Southern African rock python (*Python natalensis*), black mamba (*Dendroaspis polylepis*), puff adder (*Bitis arietans*), Mozambique spitting cobra (*Naja mossambica*) and the spotted bush snake (*Philothamnus semivariegatus*). Large rock monitors (*Varanus albigularis*) are also common and occasionally lead to alarm calling. Other reptile species that provoke alarms are the Nile crocodile (*Crocodylus niloticus*), Nile monitor (*Varanus niloticus*) and leopard tortoise (*Stigmochelys pardalis*), which were all excluded from analysis since they are only encountered very rarely.

There is no data on mortality due to snake predation for our population. However, we can report one confirmed case of an adult female from one group that was caught by a python in January 2021. The respective individual carried a radio telemetry collar at the time of predation and our observers, while searching for the group, found the python emitting the signal, with all other monkeys of the group still close by. As encounters with pythons are common at our field site, we assume that python predation is a normal cause of mortality for our vervet population. While several of the most frequently encountered snake species are venomous, including black mambas, Mozambique spitting cobras and puff adders, we have not observed any instances of deadly snake bites (but see Henzi et al. 2021).

### 2.3 | Ad libitum data collection

Groups were observed in two time shifts, either starting at sunrise or eight hours before sunset. We scored individual daily presence in group logbooks and collected ad-libitum data on natural encounters with snakes and monitor lizards (hereafter ‘reptile event’). In all reptile events, we recorded which individuals of the group approached and inspected the reptile within a 10-meter radius (inspect: yes/no), as well as which subjects of the group produced alarm calls (alarm: yes/no). We took the first alarm calls as an indicator that a reptile had been discovered and scored event duration as the approximate time that passed between the first calls and when the last subjects left the 10 m radius around the threat. As precise measurements of event durations are hard to obtain with ad libitum sampling, we classified duration on an ordinal scale of 5-min intervals. Lastly, we noted the GPS position of the event and the species that was encountered. In total, we recorded 206 reptile events where observers were present at the start of the event and could identify the reptile that the monkeys responded to. Whenever possible we roughly estimated length and diameter of snake species (Fig. S1), but did not use these estimates for statistical models as they are error prone and could not be obtained when snakes were curled.

### 2.4 | Python model presentations

We crossed the factors snake length (1 m and 3 m) with snake diameter (5 cm and 15 cm) and constructed one snake model for each of the four length/diameter combinations. All models were made from the same artificial leather imitating the scale pattern of real pythons (ESM, Fig. S2 – S4). Each group received a single presentation with each of the four snake models, in a balanced and pseudorandomized order, spaced three to four weeks apart to avoid habituation.

In each presentation, one observer placed the snake model ahead of the group (distance ≥ 100 m) into their estimated travel path, while maintaining constant radio contact with the other two observers that monitored the group to ensure that the setup occurred out of sight of the monkeys. After preparing the snake model, one observer positioned themselves, partially concealed by vegetation, parallel to the model (distance 30 - 50 m), while the other observers remained close to the most peripheral subjects of the group. Having a concealed observer wait at a distance from the model, while the other observers approached in tandem with the groups’ frontline ensured that we recorded the moment when the monkeys found the snake model, while avoiding that the monkeys would ‘discover’ the snake models in close proximity to observers. We chose this approach to avoid habituation and imitate a ‘natural’ event structure, where human observers would approach a potential snake together with the monkeys, following the calls of the individuals who found the snake. We only presented at locations that resembled natural encounter spots for pythons, to make our presentations as realistic as possible.

After the monkeys had discovered the snake model, we recorded the identity of all subjects that inspected the model within a 10-meter radius (inspect: yes/no) and produced alarm calls (alarm: yes/no), with all three observers carrying video cameras for documentation. A trial was considered to be over when the entire group had departed and was out of sight (distance ≥ 100 m). All experiments took place between March and July 2021. We only conducted experiments in five of the six groups, because the habituation state of one recently re-habituated group (CR) did not yet allow experiments.

### 2.5 | Statistical analysis – General procedure

Statistical analysis was conducted in R (v. 4.3) (R Core Team, 2023) using the brms package (v. 2.20.4) (Bürkner, 2017). We fitted an ordinal model with cumulative logit link function to analyze the duration of natural snake encounters and binomial models with a logit link function to analyze individual inspection and alarm calling probability in natural encounters and predator model presentations. We included all theoretically identifiable random slopes into each model. We z-transformed the covariates group size and trial number to a mean of zero and a standard deviation of 1 and, in the random effects parts of the models, dummy coded and centered the factors sex (female/male), age group (juvenile/adult), snake model length (1m/3m) and snake model diameter (5cm/15cm). In the analysis of natural snake encounters, we modelled reptile ID as a random effect in all models since the kind of reptiles that are most frequently observed by our population are likely not representative of the full set of possible reptile species that vervet monkeys may encounter. All models were run with four chains for 6000 iterations, specifying weakly informative priors (normal (0,1)) for the main effects. Model convergence was confirmed in all cases (R = 1.00, bulk and tail ESS > 700) and model performance was assessed via the ‘posterior predictive check’ function (pp_check). We compared full models with random effects only models using Leave-One-Out Cross-Validation (Vehtari et al., 2017). To assess how well models explained the variance in the data, we used the ‘bayes_R2’ function to determine conditional and marginal R^2^ values. Since calculation of R^2^ assumes continuous variables with equal differences among the levels of the response variable, R^2^ values for the ordinal model analyzing event duration must to be treated with caution. Inferences were made based on model comparisons with random effect only models and 95% credible intervals (CI) of posterior distributions in conjunction with ‘probability of direction’ (PD) estimates from the ‘bayestestR’ package (Makowski et al., 2019).

### 2.6 | Natural encounters – Event duration model

We examined which variables predicted how long groups remained in close proximity to reptiles (range: ≤ 5 to ≤ 90 min), via a GLMM with ‘cumulative’ family and logit link function. We used 202 out of 206 events for this analysis, as event duration was not scored in four events (N_Black mamba_ = 3, N_Spotted bush snake_ = 1). As fixed effect, we only entered the control predictor group size (mean ± s.d. = 40.3 ± 16.5; range: 14 to 77). As random effects, we included random intercepts for group ID and the encountered reptile species as well as the random slope of group size within group ID and species.

### 2.7 | Natural encounters – Inspection and alarm calling probability models

To investigate which variables predicted whether individuals in the group inspected (yes/no) a potential threat and alarm called (yes/no) in a reptile event, we used a GLMM with ‘bernoulli’ family and logit link function. In both models (inspection and alarm calling), we used a sample of 92 from the 206 total events, where the ID of all subjects that inspected and alarm called could be determined. We included the fixed effects sex (female/male), age group (juvenile/adult) and the two-way interaction between sex and age group, as well as group size (mean ± s.d. = 44.6 ± 14.7; range: 16 to 73) as a control. We entered the random intercepts of individual ID, group ID, event ID and species and included all identifiable random slopes of main effects within the random intercepts.

### 2.8 | Snake model presentations – Inspection and alarm calling probability models

We used a GLMM with ‘bernoulli’ family and logit link function to assess whether individual probability to inspect (yes/no) and alarm call (yes/no) depended on the size of snake models. As fixed effects, we included sex (female/male), age (juvenile/adult), the interaction between sex and age group, snake model length (1m/3m) and snake model diameter (5cm/15cm). We also included trial number as a control (range: 1 to 4) to assess whether habituation affected response intensity. We did not include group size, as this variable did not vary within groups at the time of our model presentations and the random factor group hence covered variation in group size between groups. As random intercepts, we included individual ID and group ID. We entered all identifiable random slopes of main effects within the random intercepts.

## 3 | Results

### 3.1 | Natural encounters – Event duration

Over the 25-month study period (2280 group observation days) we recorded 202 encounters with identified reptiles (N_Python_ = 53, N_Black mamba_ = 38, N_Mozambique spitting cobra_ = 30, N_Puff adder_ = 16, N_Spotted bush snake_ = 26, N_Rock monitor_ = 39) throughout the home ranges of all groups (N_BD_ = 69, N_NH_ = 48, N_LT_ = 42, N_AK_ = 20, N_KB_ = 18, N_CR_ = 5). Encounters appeared to be more common close to the main river, especially for pythons (Fig. 1a). Group size had no effect on event duration (PD = 52.02%) and explained comparatively little variance in the data (R^2^_Marginal_ = 0.08 ± 0.09 SE; R^2^_Conditional_ = 0.28 ± 0.05 SE). Visual inspection of the model and raw data indicated that, while for most species short events (≤ 5 min) were typical, events with pythons and black mambas exhibited higher average event duration, but were also more variable in duration (Fig. 1b, ESM Table 2).

**Fig. 1.**
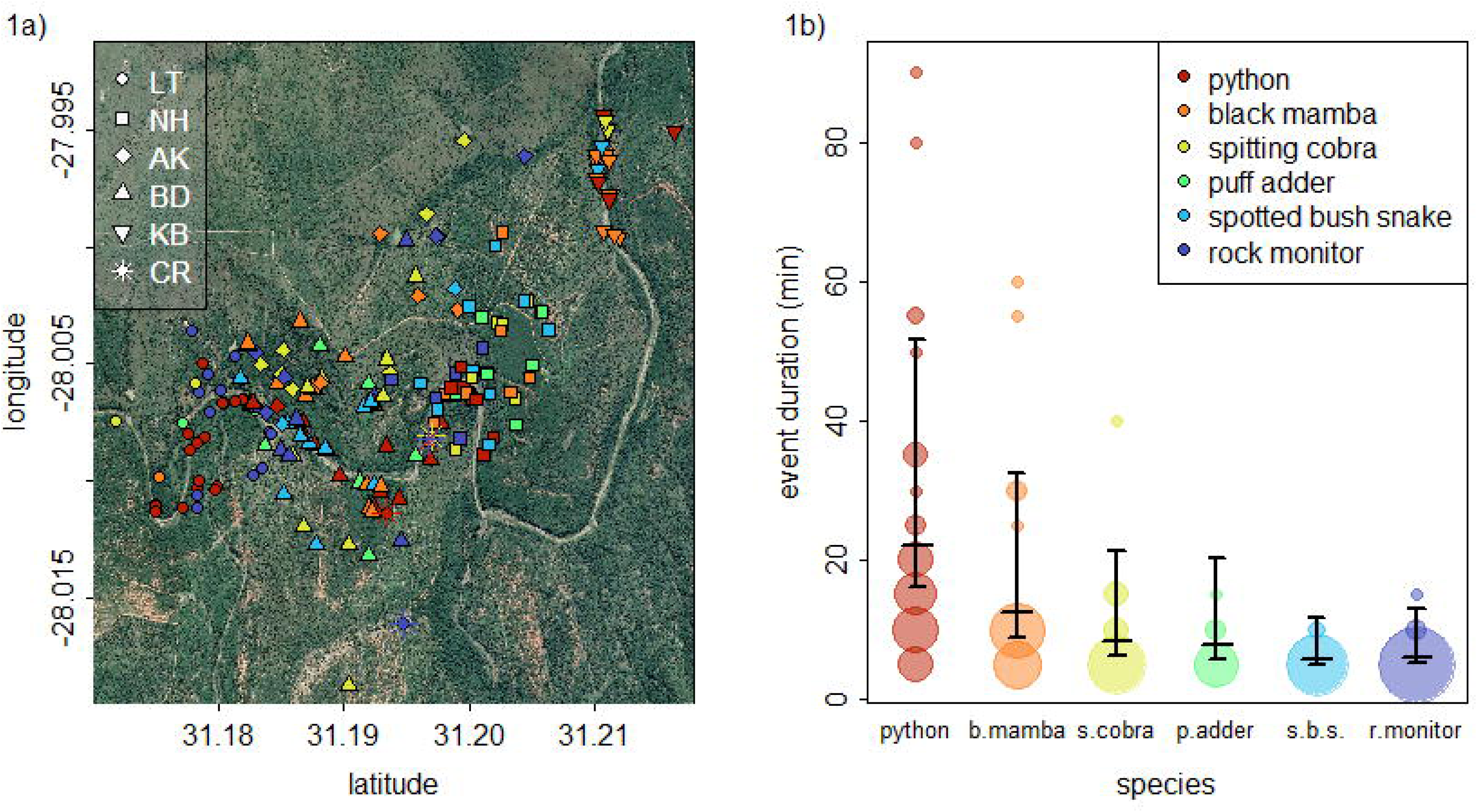
a) Location of reptile events recorded from all groups and colored by species. **b)** Variation in event duration across the different species. Points indicate the relative frequency of event durations (measured in 5 min intervals), with the size of the area corresponding to the number of observations (range: 1 to 35). Vertical lines with error bars show the posterior medians and 95% CIs for an average group size that were derived from the model by averaging event durations weighted by the predicted estimated probabilities for all event durations within species.

### 3.2 | Natural encounters – Which individuals inspected the threat?

We recorded 92 events in which all subjects that inspected and alarm called could be identified. The sample included events with all six reptile species (N_Python_ = 19, N_Black mamba_ = 19, N_Mozambique spitting cobra_ = 13, N_Puff adder_ = 11, N_Spotted bush snake_ = 19, N_Rock monitor_ = 11) and 254 individuals in five groups (N_BD_ = 25, N_NH_ = 32, N_LT_ = 16, N_AK_ = 7, N_KB_ = 12), yielding 3573 data points, with 625 inspections.

Model comparisons indicated that the full model fitted the data better than the random effect only model (model weight _full model_ = 0.81, ESM Table 3). Model estimates for all predictors exhibited wide CIs, indicating uncertainty in the magnitude of effects (Fig. 2). There was no support for an effect of group size on individual inspection probability (PD = 76.64%), but weak support for an effect of the interaction between sex and age group (PD = 98.56%), suggesting that adult males were less likely to inspect reptiles than other age and sex categories (Fig. 3a). The fixed effects explained comparatively little of the observed variation (R^2^_Marginal_ = 0.04 ± 0.04 SE) suggesting a relevant contribution of the random effects on the variance in the data (R^2^_Conditional_ = 0.28 ± 0.02 SE). Visual exploration of the model indicated that the effect of the interaction between sex and age group was consistent across the different species that were encountered (Fig. 3a). Events with pythons showed comparatively high inspection probability compared to events with all other reptiles (Fig. 3a).

**Fig. 2.**
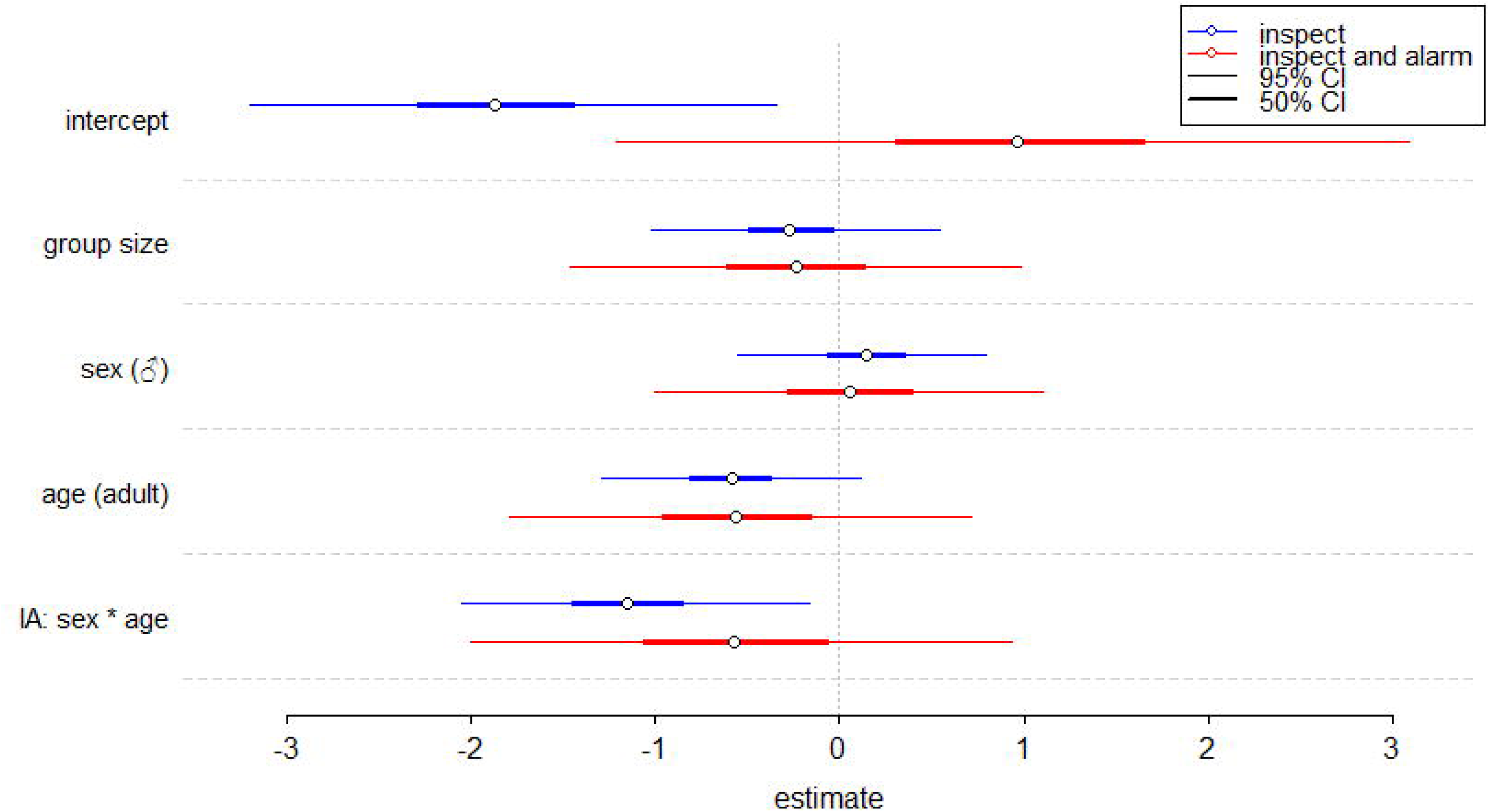
Posterior estimates for the fixed effects on inspection probability (blue) and alarm calling probability given prior inspection (red) during natural encounters of vervet monkeys with reptiles. Thick and thin lines indicate 50% and 95% CIs. The model supports an effect of the interaction between individual sex and age on inspection probability. IA: Interaction.

**Fig. 3.**
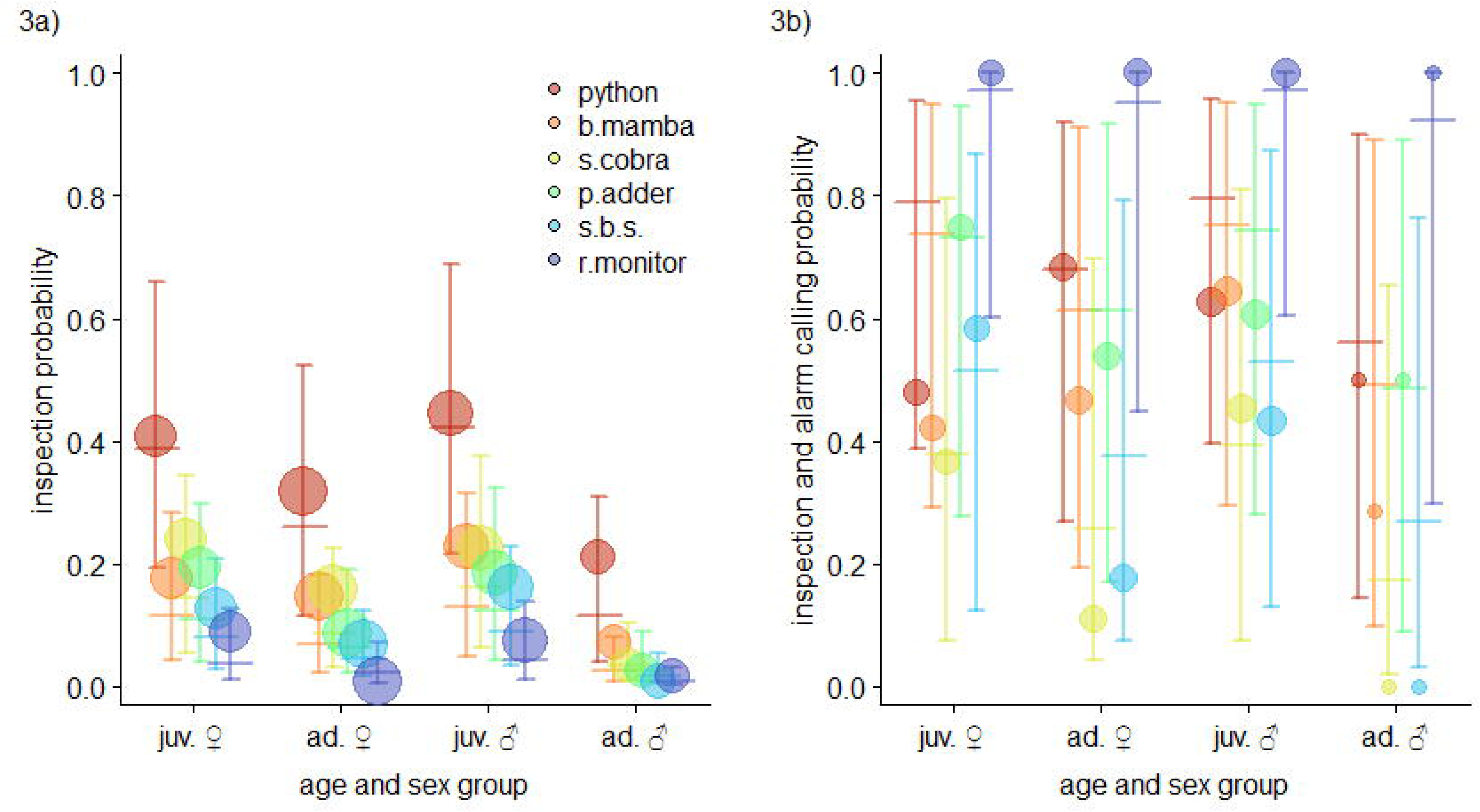
Individual inspection and alarm calling probability in natural reptile encounters relative to subject age and sex group and across all levels of the random factor reptile category. Points show the raw data average, with the area of circles corresponding to the relative frequency of observations (range 3a: 58 to 248; 3b: 1 to 91). Horizontal lines show predicted response probabilities with error bars indicating 95% CIs, assuming an average group size for **a)** inspection probability and **b)** alarm calling probability given prior inspection.

### 3.3 | Natural encounters – Which individuals alarm called?

Alarm calling was fully nested within inspection. All individuals that alarm called in an event (N_Alarms_ = 333) were also recorded as inspecting the reptile in these events, but not all individuals that inspected (N_Inspections_ = 625) also produced alarm calls. Since alarm calling was thus conditional on prior inspection, we only included those individuals into our analysis of alarm calling behavior that had been recorded as inspecting the threat to assess whether the two processes followed different rules. The reduced sample yielded 625 data points with 333 alarm calls, from 188 individuals, in 92 events and 5 groups.

The full model fitted the data only slightly better than the random effect only model (model weight _full model_ = 0.52, ESM Table 4), but wide CIs in all posterior estimates suggested large variation in the magnitude of fixed effects (Fig. 2). We found no support for an effect of group size (PD = 66.6%), sex (PD = 54.47%), age (PD = 81.77%) and the interaction between sex and age (PD = 77.46%). Fixed effects only accounted for a small degree of the variance in the data (R^2^ = 0.08 ± 0.06 SE), while the full model including random effects explained substantially more of the observed variation (R^2^_Conditional_ = 0.42 ± 0.03 SE).

We therefore explored the model visually and plotted alarm calling probability by sex and age group for all species separately (Fig. 3b). Within the set of individuals that inspected the threat, subjects appeared more likely call for pythons, black mambas, and puff adders than for spitting cobras and spotted bush snakes (Fig. 3b). While monitor lizards where only rarely inspected (Fig, 3a), subject that did inspected them always alarm called (Fig. 3b).

### 3.4 | Snake model presentations – Which individuals inspected the models?

We conducted 20 presentations in five groups gathering data from 173 individuals that provided 663 data points with 472 inspections. The random effect only model had a slightly better fit to the data than the full model (model weight _full model_ = 0.48, ESM Table 5). We found no clear support that inspection probability was affected by any of the fixed effects (Fig. 4), including sex (PD = 86.02%), age (PD = 95.28%), the interaction between sex and age (PD = 94.97%), snake model length (PD =68.43%), snake model diameter (PD = 66.36%) and trial number (PD = 95.53%). The model suggested a trend for inspection probability to decline with increasing trial number and for adults and adult males in particular to be less likely to inspect snake models in general (Fig. 4, Fig. 5a). All posterior estimates had wide CIs (Fig. 4). The random effects accounted for a substantial portion of the variance in the data (R^2^_Marginal_ = 0.14 ± 0.08 SE; R^2^_Conditional_ = 0.57 ± 0.07 SE).

**Fig. 4.**
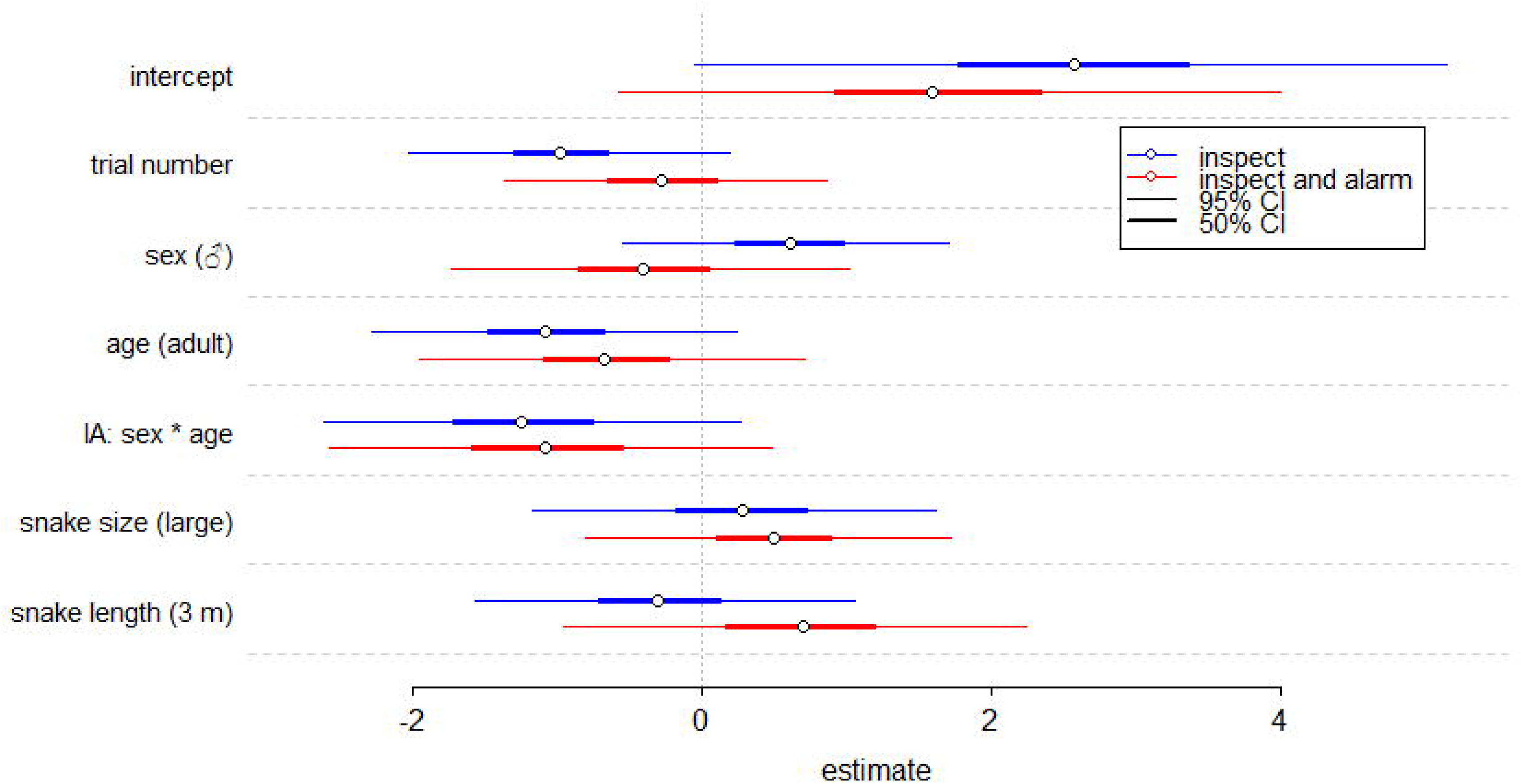
Posterior estimates for the fixed effects on inspection probability (blue) and alarm calling probability given prior inspection (red) in snake model presentations. Thick and thin lines indicate 50% and 95% CIs. The model does not provide strong support for an effect of any of the predictors. IA: Interaction.

**Fig. 5.**
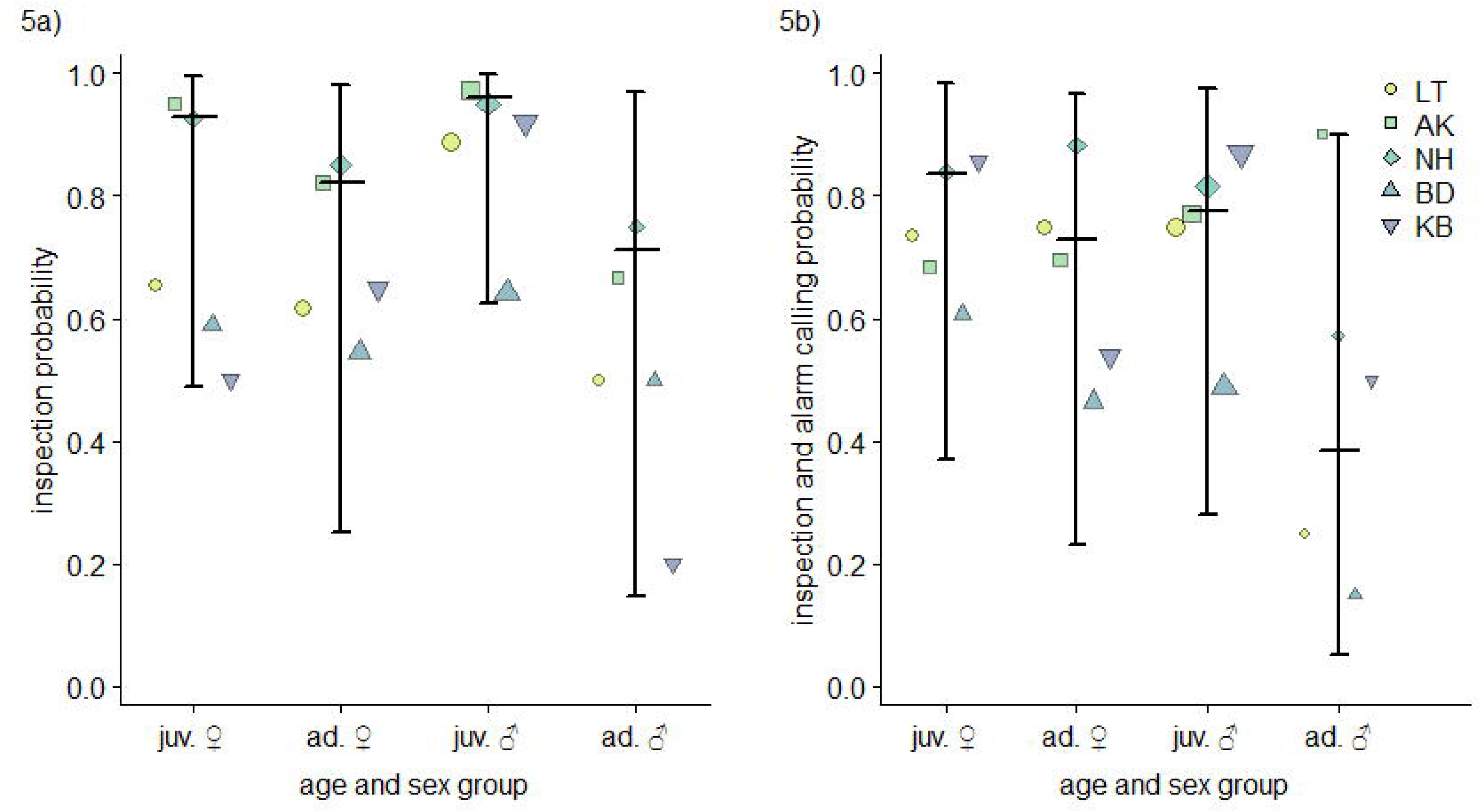
Inspection and alarm calling probability in snake model presentations relative to subject age and sex group. Symbols show the raw data average for subjects from each age and sex class pooled by group, with the area of circles corresponding to the relative frequency of observations (range 5a: 10 to 79; 5b: 2 to 43). Horizontal lines show predicted response probabilities with error bars indicating 95% CIs, assuming an average trial number, for **a)** inspection probability and **b)** alarm calling probability given prior inspection

### 3.5 | Snake model presentations – Which individuals alarm called at models?

As in our data for natural snake encounters, alarm calling in response to models (N_Alarms_ = 316) was entirely nested within inspecting models (N_Inspections_ = 472). We therefore restricted the analysis of alarm calling to those individuals that were recorded as inspecting the model, yielding a sample of 472 data points with 316 alarm calls from 161 individuals over 20 presentations and five groups. The random effect only model fitted the data better than the full model (model weight _full model_ = 0.24, ESM Table 6). None of the predictors affected individual alarm calling probability (Fig. 4; PD_Sex_ = 72.44%; PD_Age_ = 84.19%; PD_IA: Sex*Age_ = 91.18%; PD_Model length_ = 80.82%; PD_Model diameter_ = 79.9%; PD_Trial Nr._ = 69.12%). Visual exploration of the model indicated that there was a trend for alarm calling probability to decrease with trial number and for adult males to show lower alarm calling probabilities (Fig. 4, Fig. 5b). The random effects were again associated with a considerable part of the observed variation in the data (R^2^_Marginal_ = 0.12 ± 0.06 SE; R^2^ = 0.6 ± 0.08 SE).

## 4 | Discussion

Both in natural encounters and predator model presentations, the vervet monkeys’ production of alarm calls was restricted to those individuals that approached closely and inspected the snake. In most natural encounters, only a fraction of group members aggregated in close proximity to snakes and only a subset of the monkeys that inspected the snakes also produced alarm calls. The production of the ‘snake chutter’ alarm call type thus appears to be strongly associated with the visual confirmation of the presence of a threat (Crockford et al., 2012). While primates were observed to produce alarm calls when revisiting sites of recent snake encounters (van Schaik & Mitrasetia, 1990), our data suggest that seeing the potential threat is a relevant factor gating snake chutters in vervet monkeys (Fischer & Price, 2017).

Alerting conspecifics to a snake’s presence, while simultaneously recruiting them to the snake’s position, decreases their risk to fall victim to an ambush, since the snake lost the element of surprise (Carlson & Griesser, 2022; Crofoot, 2012). Snakes tend to be well concealed, requiring monkeys to be in close proximity before establishing line of sight. Since alarm calls can only serve as an effective guide if they originate close to the snake, a tendency in signalers to seek visual confirmation of a snake before producing alarms, would help receivers to localize snakes (Digweed et al., 2005). Restrictive usage of alarm calls close to the threat could thus facilitate recruitment of group members, leading to the question how signalers might benefit from recruiting.

While recruiting others towards snakes will dilute a signaler’s predation risk, it is questionable why receivers should approach snake alarm calls if doing so would put them in jeopardy (Crofoot, 2012). In our observations, snakes typically remained immobile, waiting for the monkeys to leave, or sought cover in nearby vegetation. Ending a snakes current hunting episode via perception advertisement (Carlson & Griesser, 2022; Clark, 2005; Crofoot, 2012; Curio, 1978; Gursky, 2006; Swaisgood et al., 1999) may account for the alarm call’s recruitment properties. What remains unclear in this scenario, is why an individual that discovers a snake should alert or recruit others in the first place, since evading a relatively immobile threat would suffice to avoid predation for such an individual. However, if alarm calls would decrease the predation risk of the signaler’s relatives, kin selection would explain why signalers alert and recruit nearby group members towards snakes (Wheeler, 2008), especially if costs are low.

Most natural encounters of vervet monkeys with reptiles were short and we never observed snakes to attack subjects that approached them closely, suggesting that opportunity costs of anti-snake behavior are generally low for vervets (Henzi et al., 2021). Group size had no effect on event durations but it appeared that events with pythons, the only confirmed snake predators of vervets, lasted longer and individuals of all age and sex classes were more likely to inspect them (Hernández Tienda et al., 2021; Ramakrishnan et al., 2005). Lingering longer around pythons might thus increase the chances that other group members to detect it. As in a previous experimental study (Isbell & Etting, 2017), adult males were less likely to approach and inspect snakes in natural encounters than all other age and sex groups. However, males did not show reduced alarm calling probability given that they inspected snakes. Our data thus lend mixed support to kin selection as a driver of anti-snake behavior (Phillips & Waterman, 2014; Swaisgood et al., 2003; Wheeler, 2008).

In contrast, juveniles were more likely to inspect reptiles and alarm call than adults (Corrêa & Coutinho, 1997; Ferrari & Ferrari, 1990; León et al., 2023), which is consistent with the notion that juveniles are more curious and novelty affine than adults. The increased attention juveniles direct at reptiles relative to adults might be an expression of an overall increased curiosity and excitability (Dubreuil et al., 2023; Fischer, 1998; Ramakrishnan et al., 2005), which would facilitate associative learning in the predator context (Hollén & Radford, 2009; León et al., 2023; Seyfarth & Cheney, 1980, 1986). Monitoring pythons for extended time periods could thus also allow juveniles to improve at recognizing particular shapes or scale patterns (León et al., 2023; Meno et al., 2013; Owings et al., 2001; Ramakrishnan et al., 2005).

Our data lend support to the notion that vervets modulate their response according to the risk associated with a snake species. Since pythons tend to be larger than other snake species, size might represent a robust cue for attribution of risk (Ramakrishnan et al., 2005; Swaisgood et al., 1999). However, we found no effect of snake size during our presentations of differently sized snake models. Our experiment thus indicates that vervets do not discriminate snakes solely by size, but might also consider the skin pattern.

The response intensity we observed during experiments far exceeded that of most natural python encounters, which is a cause of concern. Predator model presentations are a staple method in primatology and snake models have become a popular tool to investigate whether primates adjust their alarm calling behavior to the knowledge state of receivers (Crockford et al., 2012, 2017; Girard-Buttoz et al., 2020; León et al., 2023; Mielke et al., 2019; Schel et al., 2013). If primates respond strongly to predator models in experiments, as is the case in our presentations, the common interpretation would be that the model is sufficiently convincing and the experimental design thus suited to study predator-prey interactions. As our data show, responses towards models can exceed normal levels of anti-predator behavior, which goes unnoticed if experimental data cannot be compared to natural events. The lack of such a comparison is a common constraint, as it is difficult to collect data on actual predator events and rarely possible to observe predation directly (Corrêa & Coutinho, 1997; Gursky, 2002; Heymann, 1987; Jack et al., 2020; Perry et al., 2003; Quintino & Bicca-Marques, 2013; Ribeiro-Júnior et al., 2016; Teixeira et al., 2016).

We can only speculate as to why our vervets responded so strongly to our snake models. The only snakes that could trigger responses comparable to our experiments were pythons. However, our presentations did not elicit the full range of response intensities observed in natural python encounters and instead always provoked extreme reactions. The reason for the apparently skewed response levels might be found in the presentation regime itself. Avoiding habituation constrains the number of trials a group can receive over a certain time frame, which limits the total number of trials that can be conducted in any study relying on predator models. A low number of repetitions makes it preferable to standardize conditions between trials, but unfortunately limits generalizability of results, as conditions are no longer representative of the full spectrum of socio-ecological variation within natural snake encounters. One variable that we assume to be relevant, yet cannot meaningfully measure in the field, is how spread out a group is before an experiment. If group spread appears large, not all subjects have a similar chance to approach the snake model and group movement becomes exceedingly hard to predict, making it undesirable to conduct experiments under such conditions. We therefore suggest that the comparatively high response intensity in our experiments might be related to a potential bias towards low group spread before predator model presentations. The potential for sampling bias in predator model experiments thus deserves serious consideration when choosing response variables and interpreting results.

In conclusion, alerting and recruiting nearby kin to the position of a threat and thereby decreasing their predation risk seems to be a plausible function of vervet snake alarms. Whether alarm calling and remaining in close proximity to snakes also reduces the predation risk of signalers directly, or affects snake behavior requires further study. Our data also support the notion that vervets adjust their anti-predator behavior to the level of threat a particular snake species poses to them. If vervets discriminated snakes according to associated risk, we would expect this classification to be reflected in their alarm call’s structure and rate (McLachlan & Magrath, 2020), an idea we recommend future studies to explore further. The tendency of adult males to inspect snakes less frequently, and the potential for social learning to guide the gradual adjustment of juvenile responses to adult like levels, also require further study. Although the data from of our experiments exhibited the same trends we observed in natural snake encounters, response levels in experiments were comparatively high. We therefore must urge to consider possible sampling bias in data acquired in predator experiments and compare them to data from natural predator-prey interactions whenever possible.

## Supporting information

ESM

## Acknowledgements

We are eternally grateful to all members of the IVP field team who contributed to data collection and remained at the IVP field site despite the COVID-19 pandemic. Special thanks go to Michael Henshall and Nokubonga Dlamini who managed the field site, assisted with data collection and trained new researchers after their arrival. We are grateful to the van der Walt family, owners of Mawana Game Reserve, for giving us the permission to conduct the study on their land.

## Authors’ contributions

L.S.: conceptualization, data curation, formal analysis, funding acquisition, investigation, methodology, project administration, software, visualization, Writing – original draft and Writing – review and editing. E.v.d.W.: funding acquisition, resources, Writing – review and editing. J.F.: conceptualization, funding acquisition, resources, supervision, Writing – review and editing.

## Data availability

All data and code are available on OSF under the following link: https://osf.io/68xpf/?view_only=4d9b4dcfebf042e18e15062637b17c2b

## Ethics

This research adhered to the Association for the Study of Animal Behaviour Guidelines for the Use of Animals in Research (Behaviour, 2018), the regulations set by the local authority, Ezemvelo KZN Wildlife, as well as the Animal Care Committee at the German Primate Center.

---

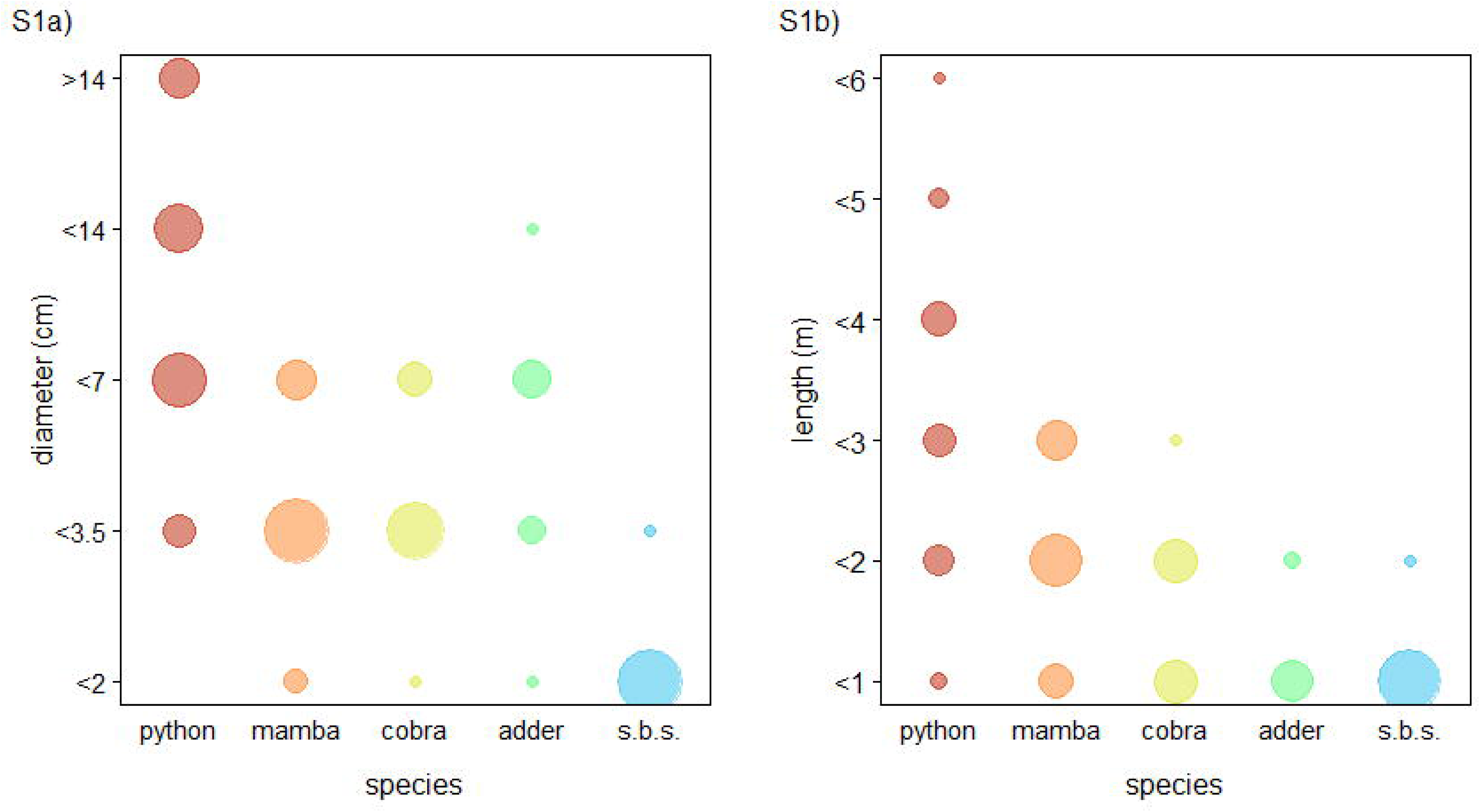

