## Supplementary material for "Anti-snake behavior and snake discrimination in vervet monkeys": ESM

**Electronic supplementary material for**

**Table 1:** Group composition (mean ± standard deviation), observation days and reptile encounters by group from April 2020 to April 2022. Data collection on the CR group was interrupted in 2020 due to the Covid-19 pandemic.

| **Group** | **BD** | **NH** | **LT** | **AK** | **KB** | **CR** |
|---|---|---|---|---|---|---|
| **Males** | 9.7 ± 2 | 6.2 ± 1.6 | 3.3 ± 1.1 | 3.1 ± 1.3 | 2.1 ± 0.9 | 3.5 ± 0.8 |
| **Females** | 19.5 ± 3.1 | 11.1 ± 2.1 | 9.2 ± 1.5 | 7.3 ± 0.8 | 4.6 ± 1.3 | 6.6 ± 2.2 |
| **Group size** | 58.8 ± 10.2 | 37.8 ± 5.8 | 26.5 ± 3.8 | 23.4 ± 3.7 | 15.9 ± 4.7 | 21.1 ± 7.8 |
| **Observation days** | 554 | 518 | 388 | 462 | 252 | 106 |
| **Total reptile events** | 71 | 48 | 42 | 20 | 20 | 5 |
| Pythons | 17 | 10 | 19 | 1 | 4 | 2 |
| Black mamba | 15 | 9 | 1 | 4 | 12 | 0 |
| Spitting cobra | 11 | 7 | 2 | 7 | 2 | 1 |
| Puff adder | 6 | 7 | 3 | 0 | 0 | 0 |
| Spotted bush snake | 12 | 9 | 1 | 3 | 2 | 0 |
| Rock monitor | 10 | 6 | 16 | 5 | 0 | 2 |

**Table 2 Model 1 results – Natural encounters – Event duration**

Model formula and specifications:

event_duration ~ z.group size +

(1 + z.group size | group) +

(1 + z.group size | species),

family = cumulative("logit"), control = list(adapt_delta=0.99), prior = prior("normal(0,1)", class = "b"), iter = 6000

Shown are model estimates, estimated errors, 95% credible intervals (CI), Rhat, and Bulk and Tail effective sample size (ESS) for fixed effects, random intercepts and slopes (sd) and correlation parameters (cor). The covariate group size was z-transformed to a mean of 0 and a standard deviation of 1.

| **fixed effects** | estimate | est. error | lower 95% CI | upper 95% CI | Rhat | bulk_ESS | tail_ESS |
|---|---|---|---|---|---|---|---|
| intercept[1] | -2.68 | 0.90 | -4.43 | -0.91 | 1.00 | 6306 | 5850 |
| intercept[2] | -1.16 | 0.87 | -2.85 | 0.55 | 1.00 | 6682 | 6127 |
| intercept[3] | -0.36 | 0.86 | -2.01 | 1.35 | 1.00 | 7041 | 6113 |
| intercept[4] | 0.12 | 0.85 | -1.51 | 1.82 | 1.00 | 7200 | 6503 |
| intercept[5] | 0.43 | 0.84 | -1.19 | 2.13 | 1.00 | 7189 | 6404 |
| intercept[6] | 0.80 | 0.84 | -0.81 | 2.52 | 1.00 | 7353 | 6772 |
| intercept[7] | 1.27 | 0.85 | -0.33 | 3.00 | 1.00 | 7435 | 7059 |
| intercept[8] | 1.50 | 0.85 | -0.10 | 3.22 | 1.00 | 7579 | 7083 |
| intercept[9] | 1.76 | 0.86 | 0.14 | 3.52 | 1.00 | 7985 | 6975 |
| intercept[10] | 2.50 | 0.92 | 0.78 | 4.42 | 1.00 | 8771 | 7659 |
| intercept[11] | 3.11 | 1.01 | 1.26 | 5.23 | 1.00 | 9654 | 7823 |
| intercept[12] | 4.22 | 1.28 | 2.01 | 7.05 | 1.00 | 12373 | 9109 |
| group size | 0.00 | 0.50 | -0.95 | 1.09 | 1.00 | 7336 | 6120 |
| **random factor: group (6 levels)** | estimate | est. error | lower 95% CI | upper 95% CI | Rhat | bulk_ESS | tail_ESS |
| sd(intercept) | 0.85 | 0.87 | 0.03 | 3.26 | 1.00 | 2690 | 3352 |
| sd(group size) | 0.66 | 0.58 | 0.03 | 2.15 | 1.00 | 4357 | 5741 |
| cor(intercept, group size) | 0.13 | 0.59 | -0.94 | 0.97 | 1.00 | 7938 | 7949 |
| **random factor: species (6 levels)** | estimate | est. error | lower 95% CI | upper 95% CI | Rhat | bulk_ESS | tail_ESS |
| sd(intercept) | 3.68 | 1.47 | 1.49 | 7.19 | 1.00 | 2832 | 3204 |
| sd(group size) | 0.40 | 0.36 | 0.01 | 1.36 | 1.00 | 5258 | 5685 |
| cor(intercept, group size) | 0.03 | 0.59 | -0.94 | 0.96 | 1.00 | 11726 | 8136 |

### Table 3 Model 2 results – Natural encounters – Inspection probability

Model formula and specifications:

inspect(y/n) ~ sex * age group + z.group size +

(1 + z.group size | individual) +

(1 + sex.1 * age group.1 + z.group size | group) +

(1 + sex.1 * age group.1 | event_ID) +

(1 + sex.1 * age group.1 + z. group size | species),

family = bernoulli(), control = list(adapt_delta=0.99), prior = prior("normal(0,1)", class = "b"), iter = 6000

Shown are model estimates, estimated errors, 95% credible intervals (CI), Rhat, and Bulk and Tail effective sample size (ESS) for fixed effects, random intercepts and slopes (sd) and correlation parameters (cor). The covariate group size was z-transformed to a mean of 0 and a standard deviation of 1. Factors (sex and age group) were dummy coded and centered in the random effects part (reference category). IA: interaction

| **fixed effects** | estimate | est. error | lower 95% CI | upper 95% CI | Rhat | bulk_ESS | tail_ESS |
|---|---|---|---|---|---|---|---|
| intercept | -1.85 | 0.72 | -3.20 | -0.35 | 1.00 | 4832 | 5914 |
| sex (male) | 0.14 | 0.34 | -0.56 | 0.80 | 1.00 | 7548 | 7670 |
| age group (adult) | -0.59 | 0.36 | -1.29 | 0.12 | 1.00 | 8495 | 8380 |
| group size | -0.26 | 0.39 | -1.02 | 0.55 | 1.00 | 7199 | 7478 |
| IA: sex * age group | -1.14 | 0.48 | -2.05 | -0.16 | 1.00 | 7958 | 8208 |
| **random factor: event ID (92 levels)** | estimate | est. error | lower 95% CI | upper 95% CI | Rhat | bulk_ESS | tail_ESS |
| sd(Intercept) | 1.07 | 0.13 | 0.85 | 1.34 | 1.00 | 4143 | 7172 |
| sd(sex) | 0.30 | 0.19 | 0.01 | 0.69 | 1.00 | 3956 | 4655 |
| sd(age group) | 0.42 | 0.21 | 0.04 | 0.82 | 1.00 | 2859 | 3788 |
| sd(ID: sex * age group) | 0.85 | 0.43 | 0.07 | 1.71 | 1.00 | 2825 | 2820 |
| cor(intercept, sex) | -0.17 | 0.37 | -0.82 | 0.62 | 1.00 | 10326 | 8465 |
| cor(intercept, age group) | -0.36 | 0.32 | -0.87 | 0.34 | 1.00 | 8724 | 7527 |
| cor(sex, age group) | 0.13 | 0.42 | -0.71 | 0.84 | 1.00 | 5320 | 7943 |
| cor(intercept, IA: sex*age group) | 0.38 | 0.32 | -0.38 | 0.86 | 1.00 | 8501 | 6909 |
| cor(sex, IA: sex*age group) | 0.03 | 0.42 | -0.77 | 0.79 | 1.00 | 6191 | 8422 |
| cor(age group, IA: sex*age group) | -0.17 | 0.40 | -0.85 | 0.64 | 1.00 | 6126 | 8952 |
| **random factor: group (5 levels)** | estimate | est. error | lower 95% CI | upper 95% CI | Rhat | bulk_ESS | tail_ESS |
| sd(intercept) | 0.51 | 0.51 | 0.02 | 1.83 | 1.00 | 5824 | 7284 |
| sd(sex) | 0.39 | 0.36 | 0.01 | 1.32 | 1.00 | 5643 | 6277 |

| | | | | | | | |
|---|---|---|---|---|---|---|---|
| sd(age group) | 0.31 | 0.32 | 0.01 | 1.14 | 1.00 | 7599 | 7889 |
| sd( group size) | 0.45 | 0.41 | 0.01 | 1.50 | 1.00 | 5566 | 5887 |
| sd(IA: sex*age group) | 0.56 | 0.53 | 0.02 | 1.94 | 1.00 | 7877 | 7209 |
| cor(intercept, sex) | -0.00 | 0.42 | -0.76 | 0.77 | 1.00 | 14700 | 9174 |
| cor(intercept, age group) | 0.02 | 0.42 | -0.76 | 0.77 | 1.00 | 18442 | 9204 |
| cor(sex, age group) | 0.01 | 0.42 | -0.75 | 0.77 | 1.00 | 13168 | 10064 |
| cor(intercept, group size) | 0.06 | 0.42 | -0.74 | 0.78 | 1.00 | 14142 | 9173 |
| cor(sex, group size) | 0.01 | 0.41 | -0.76 | 0.76 | 1.00 | 12126 | 10182 |
| cor(age group, group size) | -0.02 | 0.41 | -0.78 | 0.74 | 1.00 | 10343 | 10283 |
| cor(intercept, IA: sex* age group) | 0.02 | 0.41 | -0.75 | 0.78 | 1.00 | 19872 | 9513 |
| cor(sex, IA: sex*age group) | 0.05 | 0.41 | -0.73 | 0.79 | 1.00 | 14665 | 8947 |
| cor(age group, IA: sex*age group) | 0.02 | 0.41 | -0.75 | 0.78 | 1.00 | 10414 | 9927 |
| cor(group size, IA: sex*age group) | -0.00 | 0.41 | -0.76 | 0.76 | 1.00 | 9869 | 11023 |
| **random factor: individual (254 levels)** | estimate | est. error | lower 95% CI | upper 95% CI | Rhat | bulk_ESS | tail_ESS |
| sd(intercept) | 0.49 | 0.15 | 0.13 | 0.74 | 1.00 | 929 | 842 |
| sd(group size) | 0.36 | 0.16 | 0.05 | 0.66 | 1.00 | 890 | 2240 |
| cor(intercept, group size) | -0.43 | 0.32 | -0.94 | 0.28 | 1.00 | 2089 | 3553 |
| **random factor: species (6 levels)** | estimate | est. error | lower 95% CI | upper 95% CI | Rhat | bulk_ESS | tail_ESS |
| sd(intercept) | 1.31 | 0.57 | 0.60 | 2.76 | 1.00 | 5353 | 6823 |
| sd(sex) | 0.22 | 0.21 | 0.01 | 0.76 | 1.00 | 7936 | 6709 |
| sd(age group) | 0.41 | 0.32 | 0.02 | 1.24 | 1.00 | 5336 | 6068 |
| sd( group size) | 0.49 | 0.35 | 0.03 | 1.34 | 1.00 | 3322 | 4251 |
| sd(IA: sex*age group) | 0.49 | 0.44 | 0.02 | 1.61 | 1.00 | 6816 | 7187 |
| cor(intercept, sex) | 0.03 | 0.40 | -0.74 | 0.77 | 1.00 | 19412 | 9506 |
| cor(intercept, age group) | 0.25 | 0.38 | -0.56 | 0.86 | 1.00 | 14031 | 8397 |
| cor(sex, age group) | 0.05 | 0.41 | -0.73 | 0.79 | 1.00 | 11240 | 9687 |
| cor(intercept, group size) | 0.15 | 0.37 | -0.61 | 0.79 | 1.00 | 10832 | 7409 |
| cor(sex, group size) | 0.01 | 0.41 | -0.74 | 0.77 | 1.00 | 9281 | 8828 |
| cor(age group, group size) | 0.01 | 0.40 | -0.73 | 0.75 | 1.00 | 8070 | 8949 |

| | | | | | | | |
|---|---|---|---|---|---|---|---|
| cor(intercept, IA: sex* age group) | 0.05 | 0.4 | -0.72 | 0.77 | 1.00 | 18643 | 9123 |
| cor(sex, IA: sex*age group) | 0.07 | 0.42 | -0.72 | 0.81 | 1.00 | 12392 | 10102 |
| cor(age group, IA: sex*age group) | 0.03 | 0.41 | -0.75 | 0.78 | 1.00 | 11819 | 10396 |
| cor(group size, IA: sex*age group) | -0.01 | 0.41 | -0.76 | 0.75 | 1.00 | 10413 | 10589 |

**Table 4 Model 3 results – Natural encounter – Alarm calling probability given prior inspection**

Model formula and specifications:

alarm call(y/n) ~ sex * age group + z.group size +

(1 + z.group size | individual) +

(1 + sex.1 * age group.1 + z.group size | group) +

(1 + sex.1 * age group.1 | event_ID) +

(1 + sex.1 * age group.1 + z. group size | species),

family = bernoulli(), control = list(adapt_delta=0.99), prior = prior("normal(0,1)", class = "b"), iter = 6000

Shown are model estimates, estimated errors, 95% credible intervals (CI), Rhat, and Bulk and Tail effective sample size (ESS) for fixed effects, random intercepts and slopes (sd) and correlation parameters (cor). The covariate group size was z-transformed to a mean of 0 and a standard deviation of 1. Factors (sex and age group) were dummy coded and centered in the random effects part (reference category). IA: interaction

| **fixed effects** | estimate | est. error | lower 95% CI | upper 95% CI | Rhat | bulk_ESS | tail_ESS |
|---|---|---|---|---|---|---|---|
| intercept | 0.97 | 1.07 | -1.21 | 3.09 | 1.00 | 4324 | 5806 |
| sex (male) | 0.06 | 0.53 | -1.00 | 1.10 | 1.00 | 6869 | 8359 |
| age group (adult) | -0.56 | 0.63 | -1.79 | 0.72 | 1.00 | 7384 | 8967 |
| group size | -0.24 | 0.61 | -1.46 | 0.99 | 1.00 | 5871 | 6196 |
| IA: sex * age group | -0.56 | 0.75 | -2.00 | 0.93 | 1.00 | 8384 | 8641 |
| **random factor: event ID (92 levels)** | estimate | est. error | lower 95% CI | upper 95% CI | Rhat | bulk_ESS | tail_ESS |
| sd(Intercept) | 1.53 | 0.28 | 1.04 | 2.13 | 1.00 | 3396 | 6502 |
| sd(sex) | 0.38 | 0.29 | 0.01 | 1.10 | 1.00 | 5308 | 5293 |
| sd(age group) | 0.42 | 0.32 | 0.02 | 1.20 | 1.00 | 3996 | 4470 |
| sd(ID: sex * age group) | 1.34 | 0.89 | 0.07 | 3.31 | 1.00 | 3211 | 4265 |
| cor(intercept, sex) | 0.04 | 0.44 | -0.79 | 0.82 | 1.00 | 10677 | 7455 |
| cor(intercept, age group) | 0.06 | 0.43 | -0.76 | 0.82 | 1.00 | 10071 | 8253 |
| cor(sex, age group) | 0.09 | 0.45 | -0.77 | 0.85 | 1.00 | 7761 | 8619 |
| cor(intercept, IA: sex*age group) | 0.34 | 0.39 | -0.56 | 0.90 | 1.00 | 7216 | 7281 |
| cor(sex, IA: sex*age group) | 0.10 | 0.45 | -0.77 | 0.85 | 1.00 | 6332 | 8255 |
| cor(age group, IA: sex*age group) | -0.00 | 0.45 | -0.81 | 0.80 | 1.00 | 6405 | 8988 |
| **random factor: group (5 levels)** | estimate | est. error | lower 95% CI | upper 95% CI | Rhat | bulk_ESS | tail_ESS |
| sd(intercept) | 0.91 | 0.80 | 0.03 | 3.00 | 1.00 | 4897 | 5308 |

| | | | | | | | |
|---|---|---|---|---|---|---|---|
| sd(sex) | 0.69 | 0.59 | 0.03 | 2.18 | 1.00 | 4694 | 4779 |
| sd(age group) | 0.58 | 0.54 | 0.02 | 1.98 | 1.00 | 6022 | 5796 |
| sd( group size) | 0.98 | 0.76 | 0.04 | 2.83 | 1.00 | 3376 | 4425 |
| sd(IA: sex*age group) | 1.11 | 0.93 | 0.04 | 3.44 | 1.00 | 6126 | 5309 |
| cor(intercept, sex) | 0.01 | 0.41 | -0.75 | 0.77 | 1.00 | 9696 | 8438 |
| cor(intercept, age group) | -0.02 | 0.42 | -0.77 | 0.75 | 1.00 | 12018 | 8047 |
| cor(sex, age group) | 0.05 | 0.41 | -0.73 | 0.77 | 1.00 | 10825 | 9476 |
| cor(intercept, group size) | 0.00 | 0.41 | -0.76 | 0.75 | 1.00 | 8558 | 8853 |
| cor(sex, group size) | -0.05 | 0.41 | -0.79 | 0.74 | 1.00 | 8110 | 8504 |
| cor(age group, group size) | 0.02 | 0.41 | -0.75 | 0.77 | 1.00 | 8040 | 9157 |
| cor(intercept, IA: sex* age group) | 0.01 | 0.41 | -0.75 | 0.77 | 1.00 | 11108 | 8697 |
| cor(sex, IA: sex*age group) | -0.03 | 0.41 | -0.77 | 0.75 | 1.00 | 10458 | 8925 |
| cor(age group, IA: sex*age group) | 0.02 | 0.42 | -0.76 | 0.77 | 1.00 | 9400 | 9522 |
| cor(group size, IA: sex*age group) | 0.01 | 0.41 | -0.75 | 0.76 | 1.00 | 9278 | 10019 |
| **random factor: individual (188 levels)** | estimate | est. error | lower 95% CI | upper 95% CI | Rhat | bulk_ESS | tail_ESS |
| sd(intercept) | 0.70 | 0.33 | 0.07 | 1.34 | 1.01 | 1109 | 1675 |
| sd(group size) | 0.57 | 0.32 | 0.04 | 1.19 | 1.01 | 1120 | 2951 |
| cor(intercept, group size) | 0.35 | 0.47 | -0.79 | 0.97 | 1.00 | 1633 | 2583 |
| **random factor: species (6 levels)** | estimate | est. error | lower 95% CI | upper 95% CI | Rhat | bulk_ESS | tail_ESS |
| sd(intercept) | 1.92 | 1.04 | 0.46 | 4.50 | 1.00 | 3148 | 2657 |
| sd(sex) | 0.53 | 0.48 | 0.02 | 1.77 | 1.00 | 5942 | 5095 |
| sd(age group) | 1.32 | 0.74 | 0.26 | 3.14 | 1.00 | 4684 | 3759 |
| sd( group size) | 0.61 | 0.51 | 0.03 | 1.91 | 1.00 | 3579 | 4687 |
| sd(IA: sex*age group) | 1.38 | 0.99 | 0.07 | 3.83 | 1.00 | 5294 | 5871 |
| cor(intercept, sex) | 0.02 | 0.42 | -0.75 | 0.77 | 1.00 | 12919 | 8584 |
| cor(intercept, age group) | 0.17 | 0.38 | -0.61 | 0.83 | 1.00 | 8505 | 8853 |
| cor(sex, age group) | 0.10 | 0.41 | -0.70 | 0.81 | 1.00 | 5312 | 7761 |
| cor(intercept, group size) | 0.09 | 0.40 | -0.70 | 0.80 | 1.00 | 10894 | 8092 |
| cor(sex, group size) | 0.00 | 0.41 | -0.76 | 0.76 | 1.00 | 7513 | 8452 |

| | | | | | | | |
|---|---|---|---|---|---|---|---|
| cor(age group, group size) | 0.14 | 0.39 | -0.65 | 0.82 | 1.00 | 8302 | 9463 |
| cor(intercept, IA: sex* age group) | -0.04 | 0.40 | -0.77 | 0.72 | 1.00 | 11878 | 8265 |
| cor(sex, IA: sex*age group) | 0.02 | 0.41 | -0.75 | 0.76 | 1.00 | 7902 | 8791 |
| cor(age group, IA: sex*age group) | -0.15 | 0.39 | -0.82 | 0.63 | 1.00 | 8733 | 9001 |
| cor(group size, IA: sex*age group) | -0.06 | 0.40 | -0.78 | 0.71 | 1.00 | 8896 | 9868 |

**Table 5 Model 4 results – Snake model – Inspection probability**

Model formula and specifications:

inspection(y/n) ~ sex * age group + snake_diameter + snake_length + z.trial +

(1 + snake_diameter.1 + snake_length.1 + z.trial |individual) +

(1 + sex.1 * age group.1 + snake_diameter.1 + snake_length.1 + z.trial|group),

family = bernoulli(), control = list(adapt_delta=0.99), prior = prior("normal(0,1)", class = "b"), iter = 6000

Shown are model estimates, estimated errors, 95% credible intervals (CI), Rhat, and Bulk and Tail effective sample size (ESS) for fixed effects, random intercepts and slopes (sd) and correlation parameters (cor). The covariate trial was z-transformed to a mean of 0 and a standard deviation of 1. Factors (sex, age group, snake diameter and length) were dummy coded and centered in the random effects part (reference category). IA: interaction

| **fixed effects** | estimate | est. error | lower 95% CI | upper 95% CI | Rhat | bulk_ESS | tail_ESS |
|---|---|---|---|---|---|---|---|
| intercept | 2.57 | 1.31 | -0.05 | 5.15 | 1.00 | 4929 | 7185 |
| sex (male) | 0.61 | 0.57 | -0.55 | 1.71 | 1.00 | 11730 | 9342 |
| age group (adult) | -1.06 | 0.63 | -2.28 | 0.24 | 1.00 | 8896 | 8662 |
| snake diameter (large) | 0.27 | 0.70 | -1.16 | 1.63 | 1.00 | 11801 | 9607 |
| snake length (3m) | -0.28 | 0.66 | -1.56 | 1.06 | 1.00 | 12288 | 9190 |
| trial | -0.96 | 0.55 | -2.02 | 0.19 | 1.00 | 8258 | 8675 |
| IA: sex * age group | -1.23 | 0.73 | -2.61 | 0.27 | 1.00 | 11347 | 9398 |
| **random factor: group (5 levels)** | estimate | est. error | lower 95% CI | upper 95% CI | Rhat | bulk_ESS | tail_ESS |
| sd(intercept) | 2.46 | 1.07 | 1.08 | 5.15 | 1.00 | 4059 | 6999 |
| sd(sex) | 0.66 | 0.61 | 0.02 | 2.27 | 1.00 | 7056 | 6883 |
| sd(age group) | 1.00 | 0.84 | 0.04 | 3.13 | 1.00 | 4732 | 6502 |
| sd(snake diameter) | 1.96 | 0.99 | 0.50 | 4.37 | 1.00 | 4973 | 5086 |
| sd(snake length) | 1.06 | 0.89 | 0.04 | 3.32 | 1.00 | 6454 | 6434 |
| sd(trial) | 0.90 | 0.73 | 0.04 | 2.81 | 1.00 | 5426 | 6538 |
| sd(IA: sex*age group) | 1.93 | 1.55 | 0.08 | 5.73 | 1.00 | 4209 | 6622 |
| cor(intercept, sex) | -0.04 | 0.35 | -0.69 | 0.63 | 1.00 | 19916 | 9406 |
| cor(intercept, age group) | -0.15 | 0.34 | -0.76 | 0.56 | 1.00 | 15781 | 7705 |
| cor(sex, age group) | 0.00 | 0.35 | -0.66 | 0.68 | 1.00 | 13190 | 9403 |
| cor(intercept, snake diameter) | 0.24 | 0.31 | -0.39 | 0.77 | 1.00 | 15205 | 8673 |
| cor(sex, snake diameter) | 0.00 | 0.34 | -0.64 | 0.65 | 1.00 | 10715 | 10482 |
| cor(age group, snake | -0.14 | 0.34 | -0.75 | 0.56 | 1.00 | 9822 | 9512 |

| | | | | | | | |
|---|---|---|---|---|---|---|---|
| diameter) | | | | | | | |
| cor(intercept, snake length) | 0.00 | 0.35 | -0.65 | 0.66 | 1.00 | 20483 | 8918 |
| cor(sex, snake length) | 0.00 | 0.36 | -0.66 | 0.67 | 1.00 | 14442 | 9506 |
| cor(age group, snake length) | -0.01 | 0.35 | -0.67 | 0.65 | 1.00 | 12789 | 9503 |
| cor(snake diameter, snake length) | 0.02 | 0.35 | -0.65 | 0.68 | 1.00 | 12042 | 10118 |
| cor(intercept, trial) | -0.17 | 0.34 | -0.76 | 0.52 | 1.00 | 17670 | 9283 |
| cor(sex, trial) | 0.01 | 0.36 | -0.66 | 0.68 | 1.00 | 14423 | 9309 |
| cor(age group, trial) | 0.09 | 0.35 | -0.60 | 0.72 | 1.00 | 11691 | 10214 |
| cor(snake diameter, trial) | -0.11 | 0.34 | -0.73 | 0.58 | 1.00 | 11565 | 9816 |
| cor(snake length, trial) | 0.02 | 0.35 | -0.65 | 0.68 | 1.00 | 9639 | 10397 |
| cor(intercept, IA: sex* age group) | -0.02 | 0.34 | -0.66 | 0.62 | 1.00 | 16805 | 9382 |
| cor(sex, IA: sex*age group) | -0.04 | 0.35 | -0.70 | 0.64 | 1.00 | 11600 | 9365 |
| cor(age group, IA: sex*age group) | 0.09 | 0.35 | -0.61 | 0.72 | 1.00 | 11141 | 9975 |
| cor(snake diameter, IA: sex*age group) | -0.09 | 0.34 | -0.70 | 0.57 | 1.00 | 11321 | 10318 |
| cor(snake length, IA: sex*age group) | -0.03 | 0.35 | -0.69 | 0.64 | 1.00 | 8949 | 10358 |
| cor(trial, IA: sex*age group) | 0.06 | 0.35 | -0.63 | 0.70 | 1.00 | 8698 | 10207 |
| **random factor: individual (173 levels)** | estimate | est. error | lower 95% CI | upper 95% CI | Rhat | bulk_ESS | tail_ESS |
| sd(intercept) | 2.15 | 0.57 | 1.26 | 3.49 | 1.00 | 1345 | 3474 |
| sd(snake diameter) | 2.03 | 1.15 | 0.13 | 4.46 | 1.00 | 937 | 2226 |
| sd(snake length) | 1.10 | 0.86 | 0.04 | 3.20 | 1.00 | 1311 | 1890 |
| sd(trial) | 1.81 | 0.60 | 0.77 | 3.13 | 1.00 | 1094 | 1470 |
| cor(intercept, snake diameter) | 0.01 | 0.33 | -0.67 | 0.64 | 1.00 | 4550 | 5151 |
| cor(intercept, snake length) | 0.11 | 0.41 | -0.71 | 0.82 | 1.00 | 7614 | 8198 |
| cor(snake diameter, snake length) | -0.19 | 0.42 | -0.86 | 0.71 | 1.00 | 5425 | 7958 |
| cor(intercept, trial) | -0.33 | 0.23 | -0.74 | 0.16 | 1.00 | 3160 | 5053 |
| cor(snake diameter, trial) | -0.11 | 0.31 | -0.68 | 0.61 | 1.01 | 1348 | 1799 |
| cor(snake length, trial) | 0.08 | 0.38 | -0.70 | 0.75 | 1.00 | 772 | 1981 |

**Table 6 Model 5 results – Snake model – Alarm calling probability given prior inspection**

Model formula and specifications:

Alarm call(y/n) ~ sex * age group + snake_diameter + snake_length + z.trial +

(1 + snake_diameter.1 + snake_length.1 + z.trial |individual) +

(1 + sex.1 * age group.1 + snake_diameter.1 + snake_length.1 + z.trial|group),

family = bernoulli(), control = list(adapt_delta=0.99), prior = prior("normal(0,1)", class = "b"), iter = 6000

Shown are model estimates, estimated errors, 95% credible intervals (CI), Rhat, and Bulk and Tail effective sample size (ESS) for fixed effects, random intercepts and slopes (sd) and correlation parameters (cor). The covariate trial was z-transformed to a mean of 0 and a standard deviation of 1. Factors (sex, age group, snake diameter and length) were dummy coded and centered in the random effects part (reference category). IA: interaction

| **fixed effects** | estimate | est. error | lower 95% CI | upper 95% CI | Rhat | bulk_ESS | tail_ESS |
|---|---|---|---|---|---|---|---|
| intercept | 1.64 | 1.15 | -0.57 | 4.01 | 1.00 | 4878 | 5448 |
| sex (male) | -0.39 | 0.69 | -1.73 | 1.02 | 1.00 | 9572 | 8978 |
| age group (adult) | -0.65 | 0.67 | -1.95 | 0.72 | 1.00 | 8615 | 9216 |
| snake diameter (large) | 0.50 | 0.63 | -0.80 | 1.73 | 1.00 | 8446 | 8339 |
| snake length (3m) | 0.68 | 0.81 | -0.95 | 2.25 | 1.00 | 10393 | 8869 |
| trial | -0.26 | 0.57 | -1.35 | 0.87 | 1.00 | 6344 | 8453 |
| IA: sex * age group | -1.07 | 0.78 | -2.57 | 0.48 | 1.00 | 10647 | 9523 |
| **random factor: group (5 levels)** | estimate | est. error | lower 95% CI | upper 95% CI | Rhat | bulk_ESS | tail_ESS |
| sd(intercept) | 1.88 | 0.93 | 0.70 | 4.19 | 1.00 | 3457 | 6942 |
| sd(sex) | 1.33 | 1.02 | 0.06 | 3.91 | 1.00 | 3545 | 5451 |
| sd(age group) | 1.14 | 0.97 | 0.04 | 3.62 | 1.00 | 3828 | 6418 |
| sd(snake diameter) | 1.10 | 0.96 | 0.04 | 3.53 | 1.00 | 4502 | 6769 |
| sd(snake length) | 2.27 | 1.52 | 0.17 | 5.96 | 1.00 | 3575 | 5345 |
| sd(trial) | 0.75 | 0.65 | 0.03 | 2.40 | 1.00 | 6622 | 6553 |
| sd(IA: sex*age group) | 1.44 | 1.21 | 0.05 | 4.52 | 1.00 | 5166 | 6041 |
| cor(intercept, sex) | 0.07 | 0.34 | -0.59 | 0.69 | 1.00 | 14175 | 8990 |
| cor(intercept, age group) | 0.03 | 0.34 | -0.62 | 0.67 | 1.00 | 14028 | 8627 |
| cor(sex, age group) | 0.04 | 0.35 | -0.64 | 0.70 | 1.00 | 11806 | 9024 |
| cor(intercept, snake diameter) | -0.07 | 0.35 | -0.70 | 0.61 | 1.00 | 13241 | 9006 |
| cor(sex, snake diameter) | 0.04 | 0.36 | -0.64 | 0.70 | 1.00 | 12642 | 9709 |

| | | | | | | | |
|---|---|---|---|---|---|---|---|
| cor(age group, snake diameter) | -0.05 | 0.36 | -0.70 | 0.64 | 1.00 | 10675 | 9656 |
| cor(intercept, snake length) | 0.08 | 0.33 | -0.56 | 0.68 | 1.00 | 11483 | 8677 |
| cor(sex, snake length) | 0.05 | 0.35 | -0.62 | 0.70 | 1.00 | 10148 | 9291 |
| cor(age group, snake length) | -0.08 | 0.35 | -0.71 | 0.61 | 1.00 | 8736 | 9124 |
| cor(snake diameter, snake length) | 0.03 | 0.35 | -0.65 | 0.69 | 1.00 | 8937 | 9254 |
| cor(intercept, trial) | 0.04 | 0.35 | -0.64 | 0.70 | 1.00 | 15053 | 8997 |
| cor(sex, trial) | 0.04 | 0.35 | -0.63 | 0.68 | 1.00 | 12917 | 8730 |
| cor(age group, trial) | -0.01 | 0.35 | -0.67 | 0.67 | 1.00 | 11168 | 8157 |
| cor(snake diameter, trial) | -0.02 | 0.36 | -0.68 | 0.66 | 1.00 | 10473 | 10486 |
| cor(snake length, trial) | 0.09 | 0.35 | -0.60 | 0.73 | 1.00 | 9356 | 10549 |
| cor(intercept, IA: sex* age group) | -0.03 | 0.34 | -0.67 | 0.63 | 1.00 | 17937 | 9551 |
| cor(sex, IA: sex*age group) | 0.06 | 0.36 | -0.64 | 0.71 | 1.00 | 12054 | 9272 |
| cor(age group, IA: sex*age group) | 0.03 | 0.36 | -0.65 | 0.69 | 1.00 | 11588 | 9602 |
| cor(snake diameter, IA: sex*age group) | 0.02 | 0.36 | -0.65 | 0.69 | 1.00 | 9269 | 9151 |
| cor(snake length, IA: sex*age group) | -0.01 | 0.35 | -0.67 | 0.66 | 1.00 | 9836 | 10511 |
| cor(trial, IA: sex*age group) | 0.01 | 0.36 | -0.67 | 0.68 | 1.00 | 8238 | 10355 |
| **random factor: individual (173 levels)** | estimate | est. error | lower 95% CI | upper 95% CI | Rhat | bulk_ESS | tail_ESS |
| sd(intercept) | 2.74 | 0.78 | 1.51 | 4.57 | 1.00 | 1176 | 1969 |
| sd(snake diameter) | 2.95 | 1.20 | 0.81 | 5.59 | 1.00 | 1104 | 1148 |
| sd(snake length) | 3.47 | 1.35 | 1.05 | 6.42 | 1.00 | 1035 | 1368 |
| sd(trial) | 0.45 | 0.37 | 0.02 | 1.39 | 1.00 | 2611 | 3941 |
| cor(intercept, snake diameter) | -0.27 | 0.27 | -0.76 | 0.30 | 1.00 | 3100 | 4014 |
| cor(intercept, snake length) | 0.36 | 0.26 | -0.21 | 0.80 | 1.00 | 2987 | 4733 |
| cor(snake diameter, snake length) | -0.63 | 0.22 | -0.94 | -0.09 | 1.00 | 2200 | 2337 |
| cor(intercept, trial) | -0.10 | 0.44 | -0.85 | 0.76 | 1.00 | 9456 | 8850 |
| cor(snake diameter, trial) | 0.06 | 0.43 | -0.77 | 0.82 | 1.00 | 8775 | 8602 |
| cor(snake length, trial) | -0.04 | 0.42 | -0.80 | 0.76 | 1.00 | 10363 | 9969 |

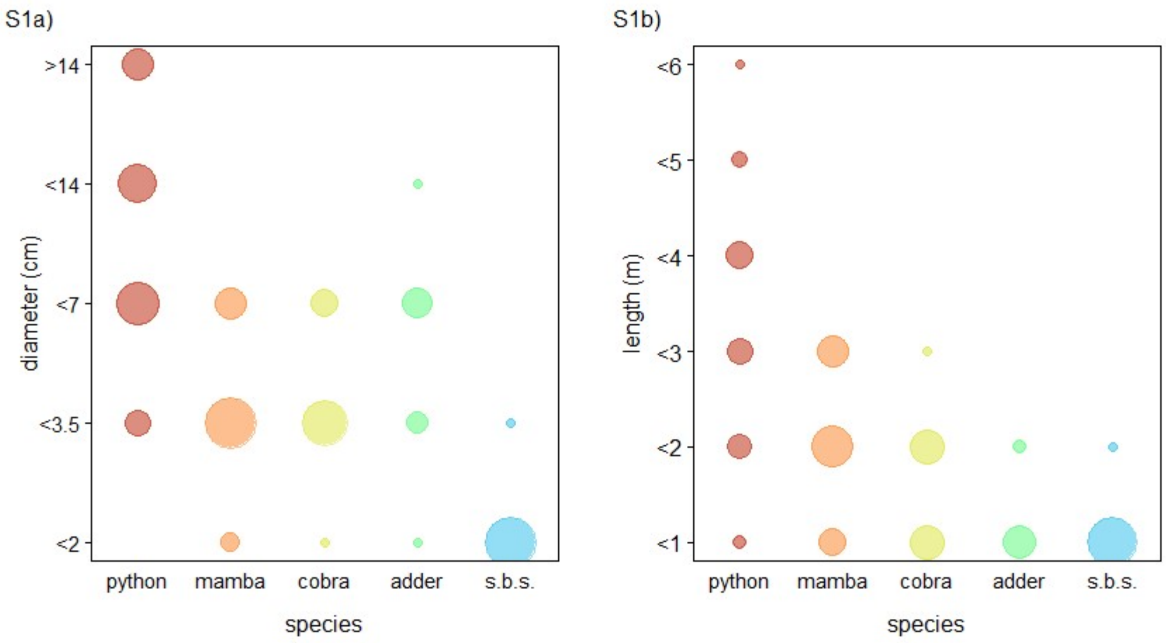


**Fig. S1** Rough estimates of a) length and b) diameter for specimen encountered in natural snake events. Diameter was estimated at the widest section of the snake (excluding the head). Area of circles is proportional to the number of observations (Fig. S1a: range = 1 to 26, N = 164; Fig. S1b: range = 1 to 25, N = 126).

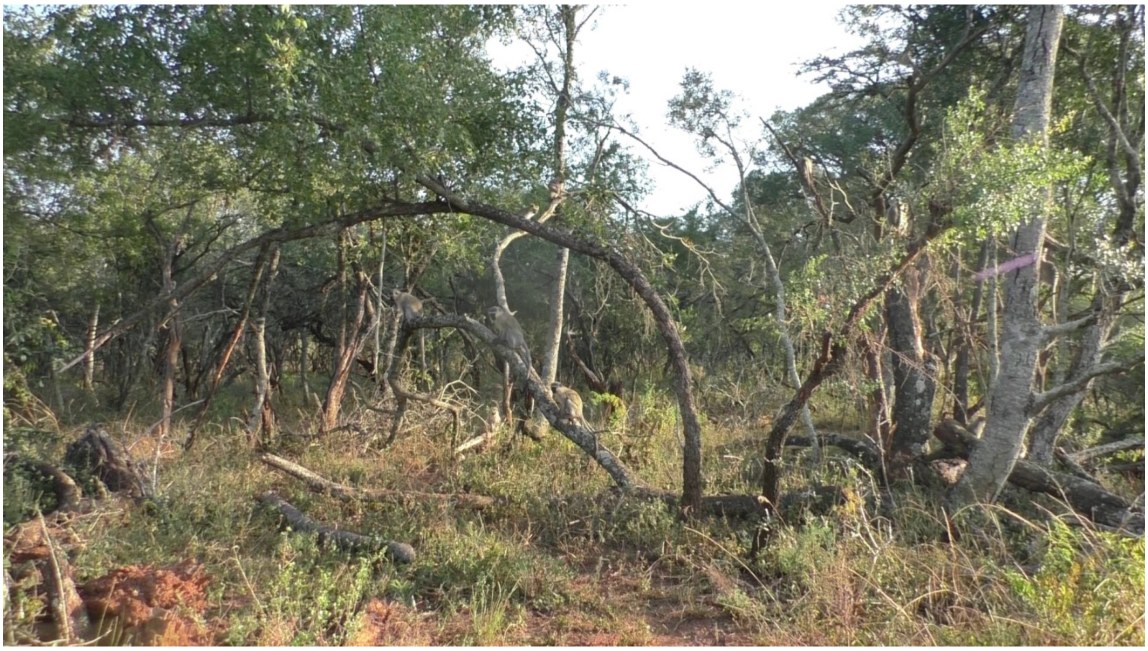

**Fig. S2** Image of snake model taken just after the first subjects arrived at the start of the presentation. Snake model in bottom left corner.

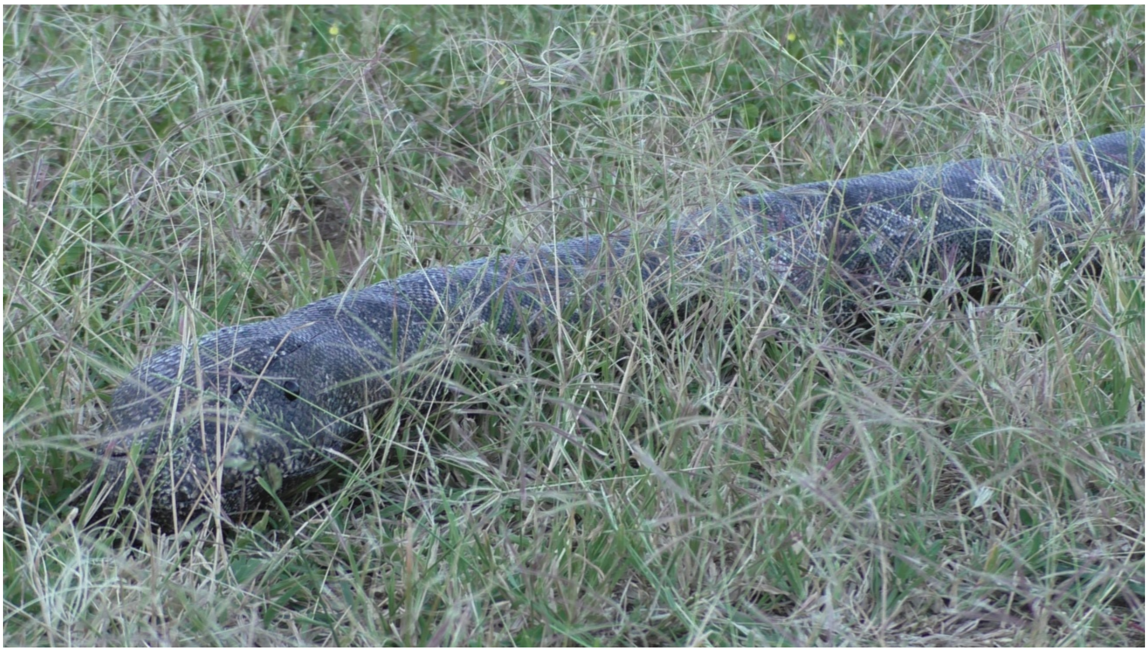

**Fig. S3** Close up image of snake model from Fig. S2.

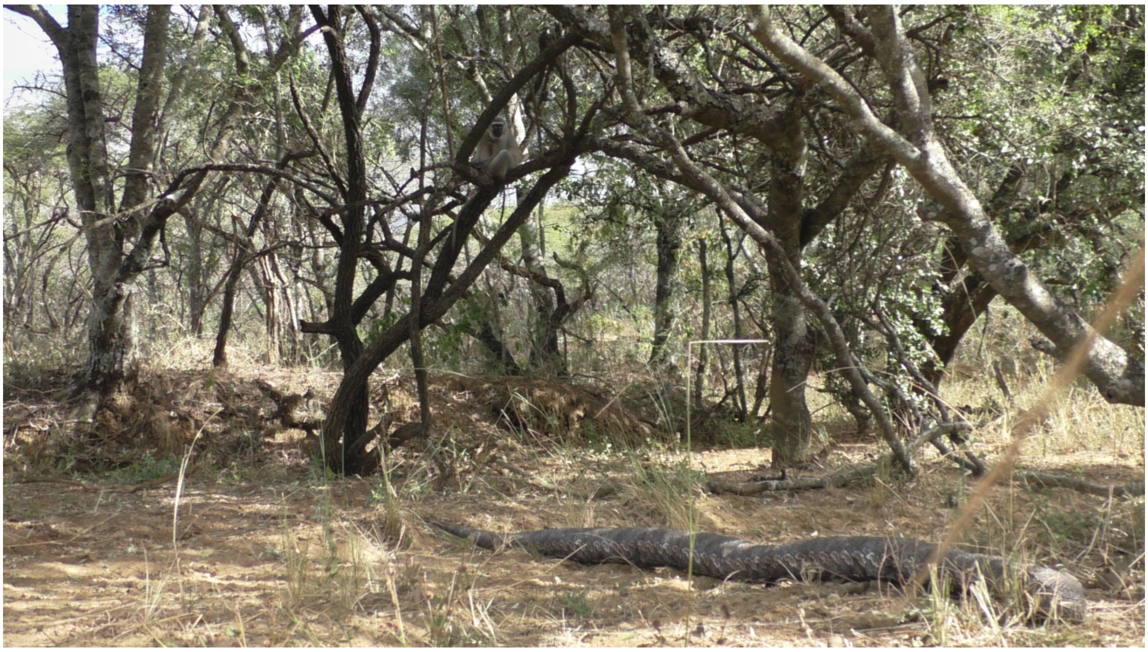

**Fig. S4** Image of snake model taken at the end of a presentation, when most subjects had left.
